# Photoreceptor-specific scene statistics reveal melanopic structure in natural environments

**DOI:** 10.1101/2025.11.10.687567

**Authors:** Niloufar Tabandeh, Manuel Spitschan

## Abstract

Natural scenes shape visual perception and non-visual responses, yet the melanopsin pathway, the retina’s key circadian input, is rarely quantified in this context. We measured spectral and spatial properties of 671 natural scenes: indoor scenes without windows (n=115), indoor scenes with windows (n=194), and outdoor scenes (n=362), across all five human photoreceptors. Radiance and photometric metrics increased systematically from windowless indoor view (mean melanopic EDI 280 lx) to indoor scenes with window view (2386 lx) to outdoor scenes (12,142 lx), with melanopic and photopic illuminance strongly correlated (*r ≈* 0.9*, p <* 10*^−^*^5^). Beyond overall *α*-opic exposure, melanopic input exhibited structured spatial variation. RMS contrast ranged from 0.001 to 0.365, lowest without windows and highest outdoors, and correlated with melanopic luminance indoors with windows (r = 0.495) and outdoors (r = 0.477) but not windowless scenes (r = 0.026). Melanopic amplitude spectra followed 1*/f* -like power laws, steeper slopes indoors (modal *≈ −*1.4) and flatter outdoors (modal *≈ −*1.1), revealing photoreceptor-resolved regularities in natural melanopic input.

## Introduction

Natural scenes are complex, and their spectral and spatial patterns shape both visual processing and broader brain functions. From an evolutionary perspective, mammalian vision has adapted to the statistical regularities of natural environments, optimising neural coding for the structure of the natural world^1–4^. Disruption of this adaptation, such as reduced exposure to spatially rich environments, has been linked to visual disorders like myopia^5^. Because the properties of natural scenes influence perceptual efficiency, visual development, and environmental adaptations, researchers have examined natural scenes from different perspectives^6–11^. Previous studies have largely focused on specific scene types, such as controlled indoor or specific types of outdoor conditions^2,12–19^, often limited to a subset of photoreceptors^19–22^. However, understanding how different environmental settings impact all five human photoreceptors (L, M, S cones, rods, and melanopsin-expressing intrinsically photosensitive retinal ganglion cells (ipRGCs)), abbreviated as *α*-opic signals^23,24^, requires a more comprehensive approach. Recent findings show that ipRGCs play a crucial role in non-visual photoreception, mediating physiological responses such as circadian entrainment, pupillary light reflexes, and regulation of alertness^25–29^. Beyond these non-visual functions, ipRGCs also modulate aspects of visual processing, including brightness perception, contrast sensitivity, and spatial integration within the visual pathway^30–36^. There is growing evidence that the intensity and spectral composition of environmental light influence human physiology and well-being through their effects on ipRGC-mediated pathways^37–39^. Although initial studies have examined how spatial features of natural scenes modulate ipRGC activation^40,41^, the mechanisms linking the spatial statistics of natural environments to *α*-opic responses remain largely unexplored.

Here, we present a photoreceptor-resolved dataset of natural scenes and use it to identify descriptive regularities in *α*-opic irradiance, radiance, contrast, and spatial frequency structure across indoor and outdoor environments. By investigating spectral metrics, spatial contrast, and amplitude spectra, we provide detailed insights into the spatial characteristics of natural scenes and their effects on photoreceptor stimulation. Special attention is given to melanopsin, a photoreceptor that plays a key role in non-visual responses like circadian rhythm regulation. We compare image-based and spectral data to explore how they complement each other in understanding natural scene statistics. Furthermore, quantifying input across all five human photoreceptors, including melanopsin, allows for more comprehensive and biologically relevant insights. These analyses reflect the real-world environments we experience through retinal input.

## Results

### A comprehensive ***α***-opic dataset of natural scenes

We introduce a new dataset of natural scenes with spectral irradiance and photoreceptor-specific radiance measurements, sampled across diverse indoor and outdoor environments. We then use this dataset to characterise how *α*-opic exposure, contrast, and spatial frequency statistics vary across scene categories. We conducted measurements at different times of day and across multiple seasons to capture natural temporal variability in illumination conditions.

The dataset consists of 671 measurements, which are categorised into three distinct view categories, indoors (n=309; indoor with window view: n=194, indoor without window view: n=115) and outdoors (n=362). The selected locations covered a broad range of spatial features (9 indoor subcategories, 11 outdoor subcategories). Importantly, these scene categories were pre-selected prior to the measurement campaign as part of the protocol definition, with the explicit objective of intentionally sampling across all predefined categories rather than selecting locations opportunistically. This protocol-driven approach was designed to ensure systematic coverage of diverse natural and built environments. For example, the predefined categories included “forest” and “open field” for outdoor scenes and “classroom” and “office” for indoor scenes. These data were gathered in five locations (Tü bingen, Germany: n=304; Munich, Germany: n=71; Prague, Czech Republic: n=35; Lyon, France: n=93; Ottawa, Canada: n=168) Figure 1C.

**Figure 1:**
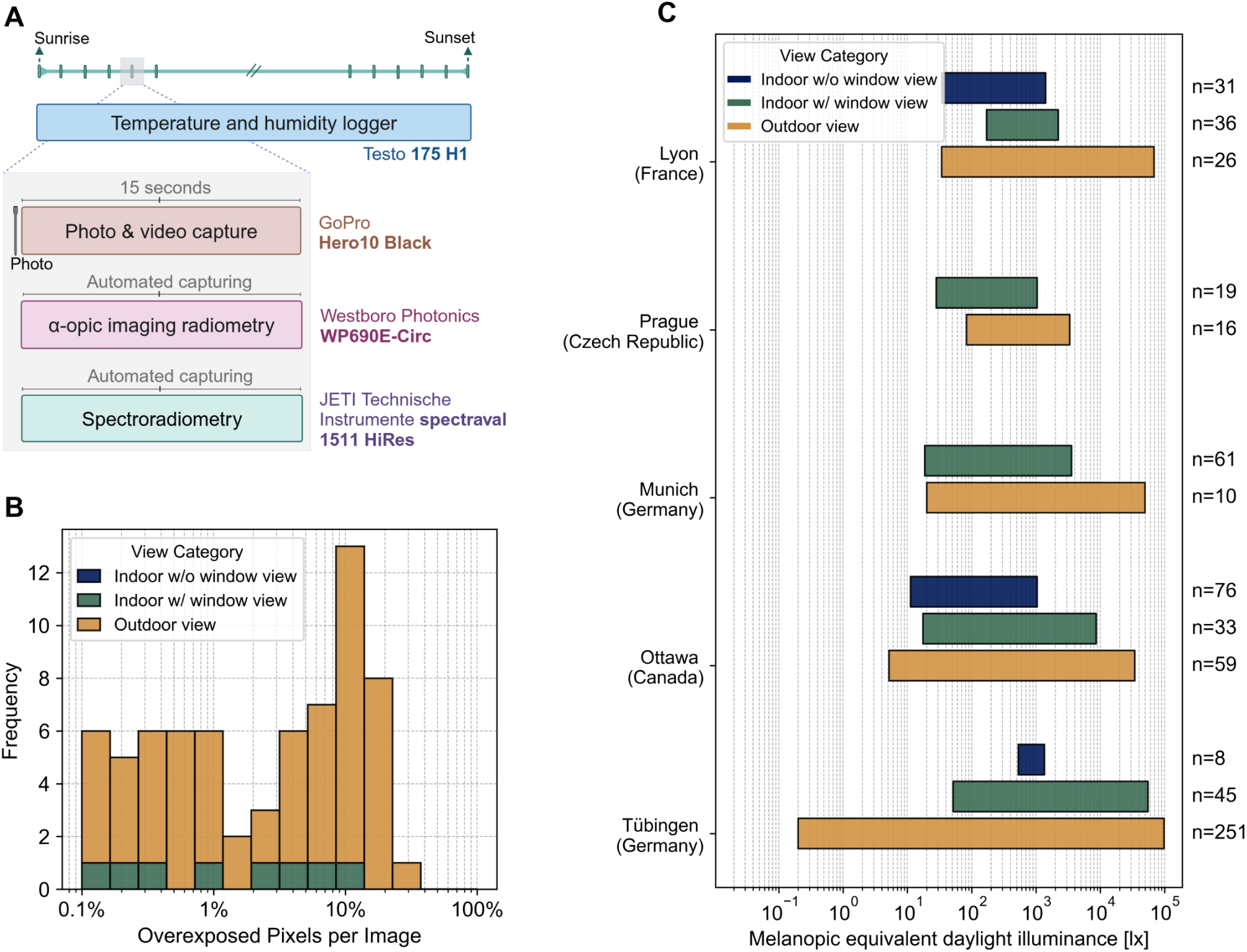
Multimodal measurements captured spectral, spatial, and environmental information across indoor and outdoor natural scenes. A.) Schematic of the measurement protocol, showing repeated measurement intervals every 30 minutes from sunrise to sunset and the devices used to capture temperature and humidity, RGB images, *α*-opic radiance images, and spectroradiometric data. B.) Stacked histogram showing the distribution of overexposed pixels in melanopsin radiance images, grouped by view category. Data are represented as image counts per logarithmic bin of percentage overexposed pixels, not as mean values or error bars. C.) Broken-bar plot showing the range of melanopic equivalent daylight illuminance [lx] across measurement locations, grouped by view category. Bars represent the minimum-to-maximum range of observed values within each location and view category, not error bars; annotated values indicate the number of measurements in each group.

The dataset includes 118 unique geometries, with additional measurements representing repeated observations of the same geometry at different times of the day. Repeated measurements were obtained for scenes in which revisiting the same geometry across different times of day was feasible and informative, allowing us to capture natural temporal variation in illumination while approximately holding spatial structure constant. An example timelapse illustrates how repeated measurements captured within-scene changes in scene appearance, photopic illuminance, melanopic EDI, and photoreceptor-specific RMS contrast across the day (see Figure S1)

To ensure the integrity of the dataset, we performed several quality checks on the collected data, focusing on missing and overexposed pixels in *α*-opic images.

#### Missing pixels

NaN values in the images are analysed across all five channels. These values represent missing pixels in a given channel and are caused by misalignment between the corresponding *α*-opic filter and lens. Consequently, each respective *α*-opic channel has the same number of missing pixels across all images in the dataset. Table 1 summarises the number and percentage of missing pixels in each *α*-opic channel. The M cone images have no missing pixels due to the perfect alignment of the M cone filter with the camera lens. The melanopsin filter shows the highest misalignment, resulting in 0.1% missing pixels in melanopsin images. However, these pixels are confined to the edges and constitute a negligible percentage of the total pixels. This ensures that they do not affect the spatial analyses and overall results.

**Table 1:** Missing pixels and their percentages across different *α*-opic channels.

|  | L cone<br>radiance map | M cone<br>radiance map | S cone<br>radiance map | Rods<br>radiance map | Melanopsin<br>radiance map |
| --- | --- | --- | --- | --- | --- |
| Missing pixels | 2712 | 0 | 6776 | 6099 | 12928 |
| Missing pixels % | 0.03 | 0 | 0.07 | 0.07 | 0.1 |

#### Overexposed pixels

Overexposed pixels occur when pixel radiance exceeds the camera’s sensor capacity to capture light intensity, resulting in an overflow of information stored as infinite (inf) values. This mostly happens due to sun glare and reflection from bright objects. We calculated the percentage of overexposed pixels for the melanopsin channel and present the distribution of images with overexposed pixel percentages above 0.1%, grouped by view category, in Table 2 and Figure 1B.

**Table 2:** Counts of melanopic radiance images with overexposed pixels across different view categories.

|  | Number of images<br>with more than 0.1%<br>overexposed pixels | Number of images<br>with more than 1%<br>overexposed pixels | Number of images<br>with more than 10%<br>overexposed pixels | Number of images<br>with more than 25%<br>overexposed pixels |
| --- | --- | --- | --- | --- |
| Indoor<br>w/o window view | 0 | 0 | 0 | 0 |
| Indoor<br>w/ window view | 8 | 5 | 1 | 0 |
| Outdoor<br>view | 61 | 36 | 19 | 1 |

Among the 675 total images, only 69 contain more than 0.1% overexposed pixels, highlighting the minimal impact of overexposure in most cases. Notably, one outdoor image exhibits overexposure exceeding 25%, underscoring the more significant variability in outdoor conditions compared to other categories. In contrast, no indoor images without a window view have overexposed pixels exceeding 0.1% of the total pixels.

### Spectroradiometric and ***α***-opic properties of natural scenes

Table 3 presents the descriptive statistics for the mean radiance values calculated for all measurements across five *α*-opic images: L cones, M cones, S cones, rods, and melanopsin. In indoor environments without a window view, radiance values are consistently low across all photoreceptor channels, with mean values ranging from 0.05 W/m²/sr (S cones) to 0.17 W/m²/sr (L cones). The small interquartile range (IQR) and low standard deviations indicate minimal variation, suggesting uniformity in pure indoor environments, although various luminaires provided lighting in different locations. In contrast, indoor scenes with a window view show a substantial increase in radiance, with mean values approximately 6 to 10 times higher than in windowless conditions.

**Table 3:** Descriptive statistics for mean radiance values across different views.

| Channel | Mean | Std | Median | Min | Max | IQR |
| --- | --- | --- | --- | --- | --- | --- |
| **Indoor without window view** |  |  |  |  |  |  |
| L cones [W/m<sup>2</sup>/sr] | 0.1687 | 0.2287 | 0.1008 | 0.0074 | 1.7264 | 0.1654 |
| M cones [W/m<sup>2</sup>/sr] | 0.1394 | 0.1670 | 0.0783 | 0.0070 | 0.8279 | 0.1211 |
| S cones [W/m<sup>2</sup>/sr] | 0.0509 | 0.0657 | 0.0209 | 0.0025 | 0.2937 | 0.0546 |
| Rods [W/m<sup>2</sup>/sr] | 0.1251 | 0.2116 | 0.0573 | 0.0061 | 1.8512 | 0.1128 |
| Melanopsin [W/m<sup>2</sup>/sr] | 0.1070 | 0.1890 | 0.0456 | 0.0050 | 1.6655 | 0.0845 |
| **Indoor with window view** |  |  |  |  |  |  |
| L cones [W/m<sup>2</sup>/sr] | 1.0201 | 1.5954 | 0.3768 | 0.0083 | 6.8467 | 0.6834 |
| M cones [W/m<sup>2</sup>/sr] | 0.9750 | 1.6437 | 0.3334 | 0.0068 | 8.2033 | 0.6957 |
| S cones [W/m<sup>2</sup>/sr] | 0.4998 | 0.8728 | 0.1526 | 0.0021 | 4.3358 | 0.3385 |
| Rods [W/m<sup>2</sup>/sr] | 0.8612 | 1.3388 | 0.3143 | 0.0048 | 5.5223 | 0.6050 |
| Melanopsin [W/m<sup>2</sup>/sr] | 0.8395 | 1.4113 | 0.2867 | 0.0036 | 6.4896 | 0.5650 |
| **Outdoor view** |  |  |  |  |  |  |
| L cones [W/m<sup>2</sup>/sr] | 2.9079 | 2.5755 | 2.1204 | 0.0004 | 10.9462 | 3.7780 |
| M cones [W/m<sup>2</sup>/sr] | 2.9978 | 3.1913 | 1.7768 | 0.0006 | 15.7464 | 3.7445 |
| S cones [W/m<sup>2</sup>/sr] | 1.6622 | 1.9117 | 0.8262 | 0.0001 | 10.2478 | 2.2489 |
| Rods [W/m<sup>2</sup>/sr] | 2.3792 | 2.1252 | 1.6200 | 0.0002 | 8.4730 | 3.2448 |
| Melanopsin [W/m<sup>2</sup>/sr] | 2.4908 | 2.5695 | 1.4308 | 0.0002 | 10.9764 | 3.2720 |

In outdoor environments, radiance levels are substantially higher, with mean values ranging between 1.6 W/m²/sr (S cones) and 2.99 W/m²/sr (M cones). The M cones, in particular, show the highest radiance levels (mean: 2.99 W/m²/sr, median: 1.77 W/m²/sr), indicating strong sensitivity to outdoor lighting conditions. This is accompanied by large standard deviations and interquartile ranges across all photoreceptor channels, reflecting the dynamic nature of natural daylight. Notably, while S cones consistently exhibit the lowest radiance values in all conditions, their increase outdoors is proportional.

Table 4 illustrates the selected derived metrics obtained from the spectroradiometer, including photopic illuminance, melanopic equivalent daylight illuminance (Melanopic EDI), and the five *α*-opic irradiance measures used in the analysis. We calculated the descriptive statistics of spectral measurements for three different views: Indoor without window view, Indoor with window view, and Outdoor view.

**Table 4:** Descriptive statistics of spectral measurements across different views.

|  | Mean | Std | Median | Min | Max | IQR |
| --- | --- | --- | --- | --- | --- | --- |
| **Indoor without window view** |  |  |  |  |  |  |
| L cone [W/m<sup>2</sup>] | 0.63 | 0.59 | 0.40 | 0.00 | 3.2 | 0.60 |
| M cone [W/m<sup>2</sup>] | 0.52 | 0.51 | 0.30 | 0.00 | 2.6 | 0.50 |
| S cone [W/m<sup>2</sup>] | 0.19 | 0.22 | 0.10 | 0.00 | 1.0 | 0.10 |
| Rods [W/m<sup>2</sup>] | 0.43 | 0.47 | 0.20 | 0.00 | 2.2 | 0.40 |
| Melanopsin [W/m<sup>2</sup>] | 0.37 | 0.42 | 0.20 | 0.00 | 1.9 | 0.30 |
| Photopic illuminance [lx] | 387.02 | 366.93 | 259.5 | 13.8 | 1959.1 | 367.00 |
| Melanopic EDI [lx] | 279.52 | 312.64 | 143.6 | 11.2 | 1416.3 | 250.50 |
| **Indoor with window view** |  |  |  |  |  |  |
| L cone [W/m<sup>2</sup>] | 4.39 | 15.09 | 1.20 | 0.00 | 103.4 | 1.575 |
| M cone [W/m<sup>2</sup>] | 3.83 | 13.05 | 1.00 | 0.00 | 89.4 | 1.40 |
| S cone [W/m<sup>2</sup>] | 1.73 | 5.64 | 0.40 | 0.00 | 38.7 | 0.70 |
| Rods [W/m<sup>2</sup>] | 3.56 | 12.07 | 0.90 | 0.00 | 82.7 | 1.20 |
| Melanopsin [W/m<sup>2</sup>] | 3.16 | 10.68 | 0.80 | 0.00 | 73.2 | 1.20 |
| Photopic illuminance [lx] | 2707.87 | 9298.21 | 723.9 | 13.9 | 63721.1 | 964.00 |
| Melanopic EDI [lx] | 2385.99 | 8050.73 | 574.75 | 17.5 | 55178.3 | 862.45 |
| **Outdoor view** |  |  |  |  |  |  |
| L cone [W/m<sup>2</sup>] | 21.63 | 34.70 | 9.40 | 0.00 | 180.8 | 16.25 |
| M cone [W/m<sup>2</sup>] | 18.76 | 29.74 | 8.45 | 0.00 | 155.1 | 14.05 |
| S cone [W/m<sup>2</sup>] | 9.39 | 13.98 | 4.60 | 0.00 | 72.7 | 7.40 |
| Rods [W/m<sup>2</sup>] | 17.89 | 27.86 | 8.35 | 0.00 | 145.2 | 13.48 |
| Melanopsin [W/m<sup>2</sup>] | 16.10 | 24.90 | 7.70 | 0.00 | 129.8 | 12.33 |
| Photopic illuminance [lx] | 13264.26 | 21277.86 | 5783.0 | 0.2 | 110872.8 | 9988.78 |
| Melanopic EDI [lx] | 12142.45 | 18765.45 | 5782.9 | 0.2 | 97807.5 | 9246.53 |

Overall, the data show a consistent trend in which the indoor with window view generally shows intermediate values between pure indoor and outdoor conditions across all derived metrics. For illuminance values (photopic illuminance and melanopic EDI) and the five *α*-opic irradiances, the outdoor view consistently exhibits the highest values, followed by the indoor with window view and the indoor without window view. This reflects higher exposure to natural light in the outdoor view, with the presence of a window modulating indoor light exposure. This pattern is more evident in the illuminance metrics. The mean photopic illuminance in the outdoor view is approximately five times higher than in the indoor with window view and 34 times higher than in the indoor without window view, with a considerably wider range of variation. The standard deviation of photopic illuminance is 21,278 lx in the outdoor view, compared to 9,298 lx in the indoor with window view and 367 lx in the indoor without window view. These results are consistent with the descriptive data from circadiometer-derived radiance images.

Notably, the variability in the indoor with window view is high across metrics, indicating a diverse range of light exposure in this setting.

### Correlational structure of ***α***-opic properties of natural scenes

Studies suggest natural scenes show consistent patterns in visual properties. Building on this, we specifically examined correlations between physiologically relevant metrics across different view categories.

Figure 2A shows the relationship between derived melanopic equivalent daylight illuminance and photopic illuminance for all measurements. As expected, melanopic equivalent daylight illuminance and photopic illuminance were strongly positively correlated across all view categories, with Pearson correlation coefficients (*r ≈* 0.9) and highly significant p-values (*p <* 10*^−^*^5^). This serves as a useful descriptive benchmark within the dataset, indicating a consistent linear association between these variables across times of day and view categories.

**Figure 2:**
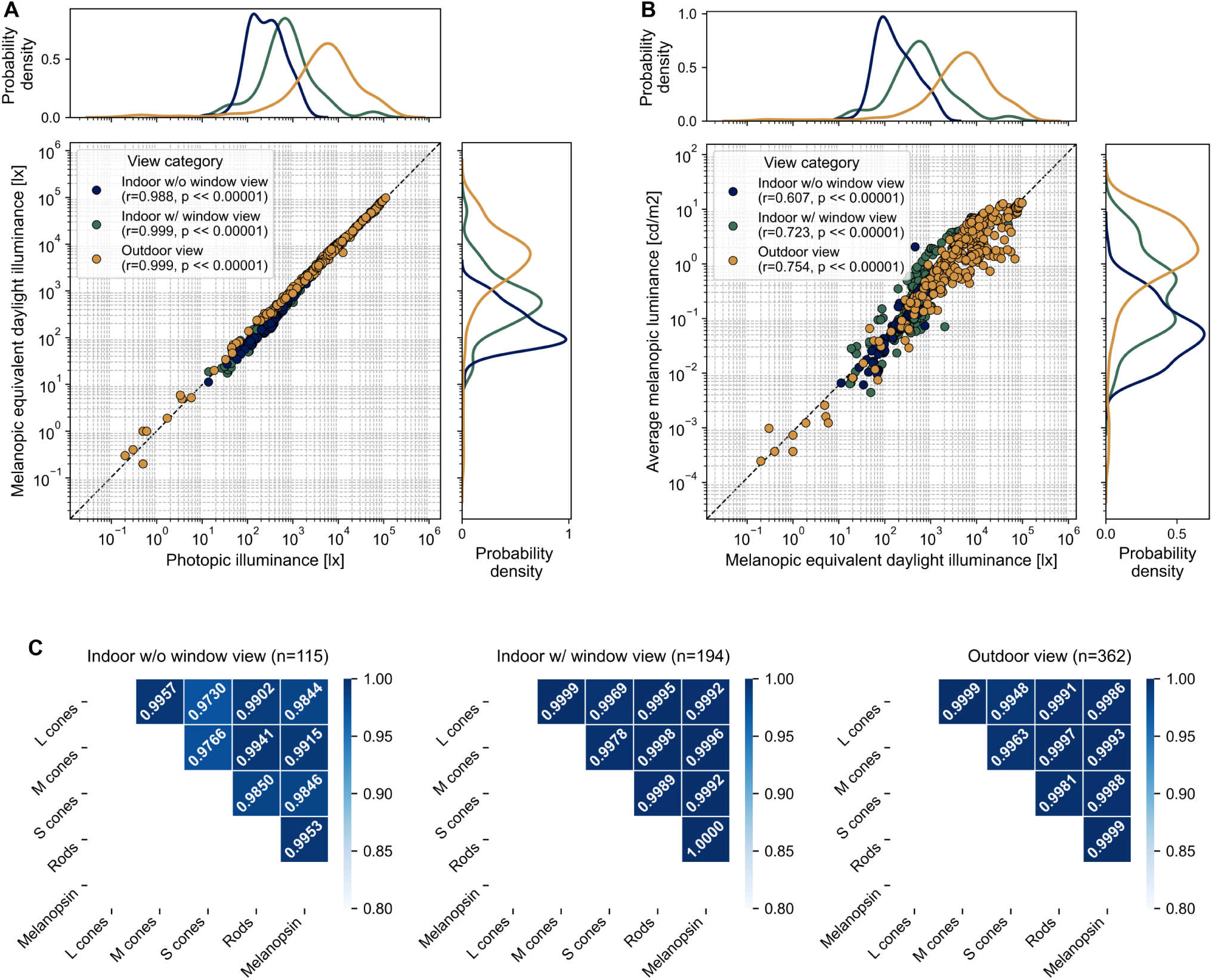
Spectroradiometric and image-derived *α*-opic measures show strong coupling across natural scenes. A.) Relationship between melanopic equivalent daylight illuminance [lx] and photopic illuminance [lx] on logarithmic axes, with marginal kernel density estimates showing the smoothed distribution of each variable. Each point represents one measurement, and data are grouped by view category. B.) Relationship between average melanopic luminance from circadiometer-derived melanopsin images and melanopic equivalent daylight illuminance from spectroradiometric measurements. Each point represents one measurement, and marginal kernel density estimates show the smoothed distribution of each variable. C.) Pearson correlation matrices for the five *α*-opic irradiance values captured by the spectroradiometer, shown separately for indoor scenes without a window view, indoor scenes with a window view, and outdoor scenes. Matrix values represent Pearson correlation coefficients.

Additionally, the probability density distributions, shown as marginal kernel density estimates (KDEs) on each scatter plot axis, reveal similar distribution patterns for both variables. The KDEs, colour-coded by view categories, illustrate that melanopic equivalent daylight illuminance and photopic illuminance distributions approximately align across different viewing conditions. This further reinforces the strong correlation observed in the main plot, as the spread and density of data points remain consistent across view categories.

The relationship between average melanopic luminance calculated from melanopic images of the circadiometer camera and melanopic equivalent daylight illuminance derived from spectral data is shown in Figure 2B. This relationship exhibits a strong overall positive correlation (*r ≈* 0.7*, p <* 10*^−^*^5^). However, the correlation varies by view category, with outdoor views showing the highest correlation (*r* = 0.754*, p <* 10*^−^*^5^) and indoor without window views having the lowest correlation (*r* = 0.607*, p <* 10*^−^*^5^).

Plotted on a logarithmic scale, the data span a broad range, with average melanopic luminance covering 10*^−^*^4^ to 10^2^ and melanopic equivalent daylight illuminance ranging from 10*^−^*^1^ to 10^6^. Despite this variation, the marginal kernel density estimates (KDEs) along each axis display similar distribution patterns across view categories and time of day. This suggests a stable relationship between spectral and image-based melanopic metrics, with outdoor environments showing a more substantial alignment between the two measures.

Figure 2C depicts the correlation matrices for the five *α*-opic irradiances derived from spectral data across three view categories. The Pearson correlation coefficients are consistently high, indicating strong linear relationships between the *α*-opic irradiances in all view categories. For example, the correlation coefficients are close to 1 for most pairs, with the lowest (*r* = 0.976*, p <* 10*^−^*^5^) between M and S cones in the indoor without window view category. This suggests a nearly perfect correlation between the *α*-opic irradiances across the different view conditions.

We also computed two sets of Pearson correlation matrices for the five photoreceptor-based radiances (L, M, S cones, rods, and melanopsin) across three view categories.

In mean radiance correlation matrices, we compute the mean radiance across all pixels for each channel and correlate these mean values between corresponding channels. As shown in Figure 3A, the Pearson correlation coefficients are consistently high across all view categories. The highest correlations are observed indoors with window conditions, where all values exceed 0.96. In contrast, the indoor without window condition shows the lowest correlations, particularly between M cones/rods (*r* = 0.7856) and M cones/melanopsin (*r* = 0.7764), indicating more significant variability in photoreceptor-relevant features in this setting.

**Figure 3:**
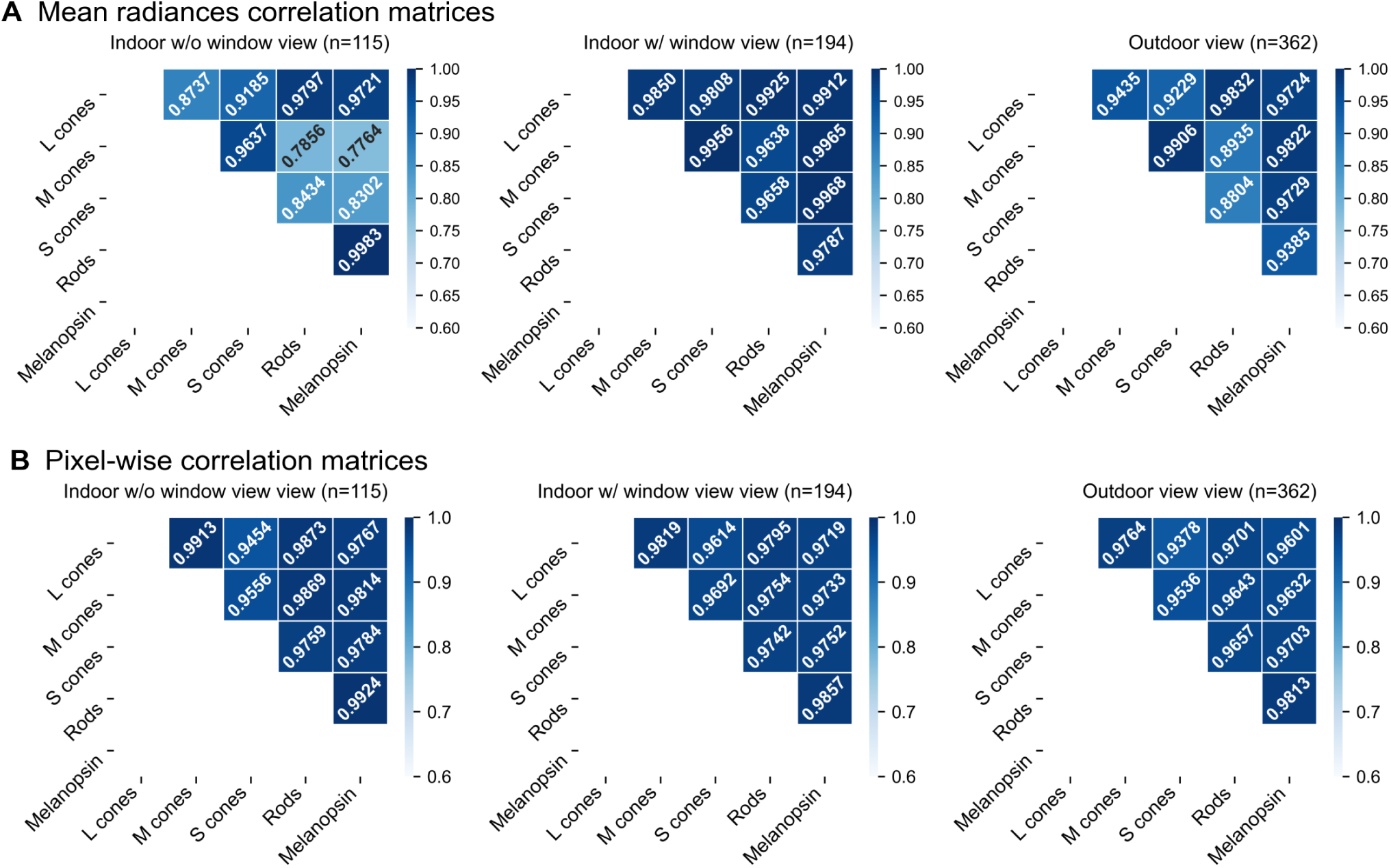
Circadiometer-derived *α*-opic radiance channels are highly correlated across view categories. A.) Pearson correlation matrices based on mean radiance values across all valid pixels in each *α*-opic radiance image, shown separately for indoor scenes without a window view, indoor scenes with a window view, and outdoor scenes. Matrix values represent Pearson correlation coefficients between *α*-opic channels. B.) Pixel-wise Pearson correlation matrices between all pairs of *α*-opic radiance channels. Correlations were computed by matching corresponding valid pixels within each image and then averaging correlation values across images within each view category. Matrix values represent the mean Pearson correlation coefficient for each channel pair.

In Figure 3B, we depict pixel-wise correlations across *α*-opic radiance images. We computed pixel-by-pixel correlations between pairs of *α*-opic channels for each image by correlating corresponding pixel values. We then averaged these correlations to obtain an overall value for each *α*-opic channel pair. Pixel-wise correlations are consistently high across all channels and views. Indoor with window and outdoor views show high correlations in both methods, but correlations appear to be stronger in the pixel-wise analysis. Indoor without window view has the lowest correlations in both cases, but the gap between channels is more pronounced in the mean-radiance correlations. Pixel-wise correlation may better preserve local correspondence, unlike the averaged method, which may smooth out important variations.

### Contrast properties of natural scenes

Spatial contrast plays a key role in visual processing, influencing how scenes are perceived and interpreted. We used the global Root mean square (RMS) contrast to quantitatively measure spatial variation in light intensity, which can differ across photoreceptor channels and scene categories. While previous studies have primarily examined contrast in luminance or colour channels^1,12,14,16,17,21,42^, this analysis extends the scope to all five human photoreceptor channels, offering a more detailed characterisation of contrast across diverse scene types.

Figure 4A reveals the RMS contrast distribution for each *α*-opic channel across view categories. As expected, the RMS contrast values of different channels show a high degree of correlation. The results show RMS contrast values ranging from 0.001 to 0.365, consistent with previous studies on spatial contrast in natural scenes^16,43^. Examples of scenes with different contrast levels are illustrated in Figure 5. The outdoor images exhibit a wider spread of contrast values compared to indoor categories, suggesting a higher variability in outdoor scenes. *Indoor with window view* images are characterised by intermediate distributions compared to the two extremes. Figure 4B shows the relationship between melanopic RMS contrast and average melanopic luminance (computed as the mean across all pixels in the melanopic channel) across three scene categories. A significant positive correlation was found for both indoor scenes with window views (*r* = 0.495, *p <* 10*^−^*^5^) and outdoor scenes (*r* = 0.477, *p <* 10*^−^*^5^). These findings suggest that scenes with higher overall melanopic luminance, such as those with daylight or outdoor views, exhibit greater melanopic RMS contrast, reflecting increased spatial variability in melanopic signal intensity. This likely reflects the influence of natural light, which introduces more variation in luminance and shadow. In contrast, indoor spaces without window views are typically lit by uniform electric lighting. As a result, no significant correlation was observed for indoor scenes without a window view (*r* = 0.026, *p* = 0.78).

**Figure 4:**
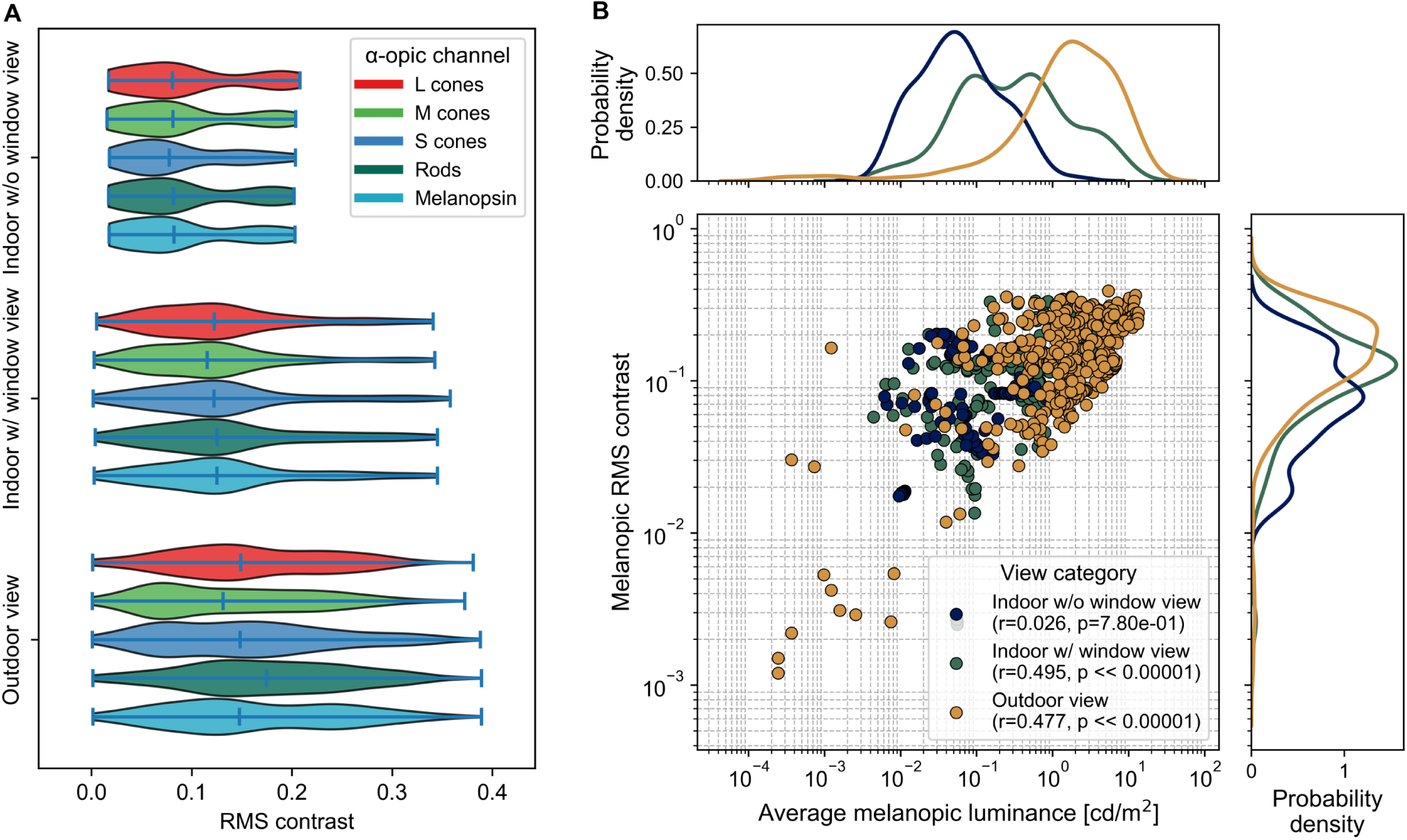
RMS contrast varies across *α*-opic channels and is related to melanopic luminance in daylight-containing scenes. A.) Violin plots showing the distribution of root mean square (RMS) contrast for each *α*-opic channel across view categories. Each violin represents the distribution of image-level RMS contrast values within the corresponding channel and view category; horizontal markers indicate medians, not means or error bars. B.) Scatter plot showing melanopic RMS contrast as a function of average melanopic luminance. Each point represents one image, and marginal kernel density estimates show the smoothed distribution of each variable. Pearson correlation coefficients and associated *p* values are shown for each view category where applicable.

**Figure 5:**
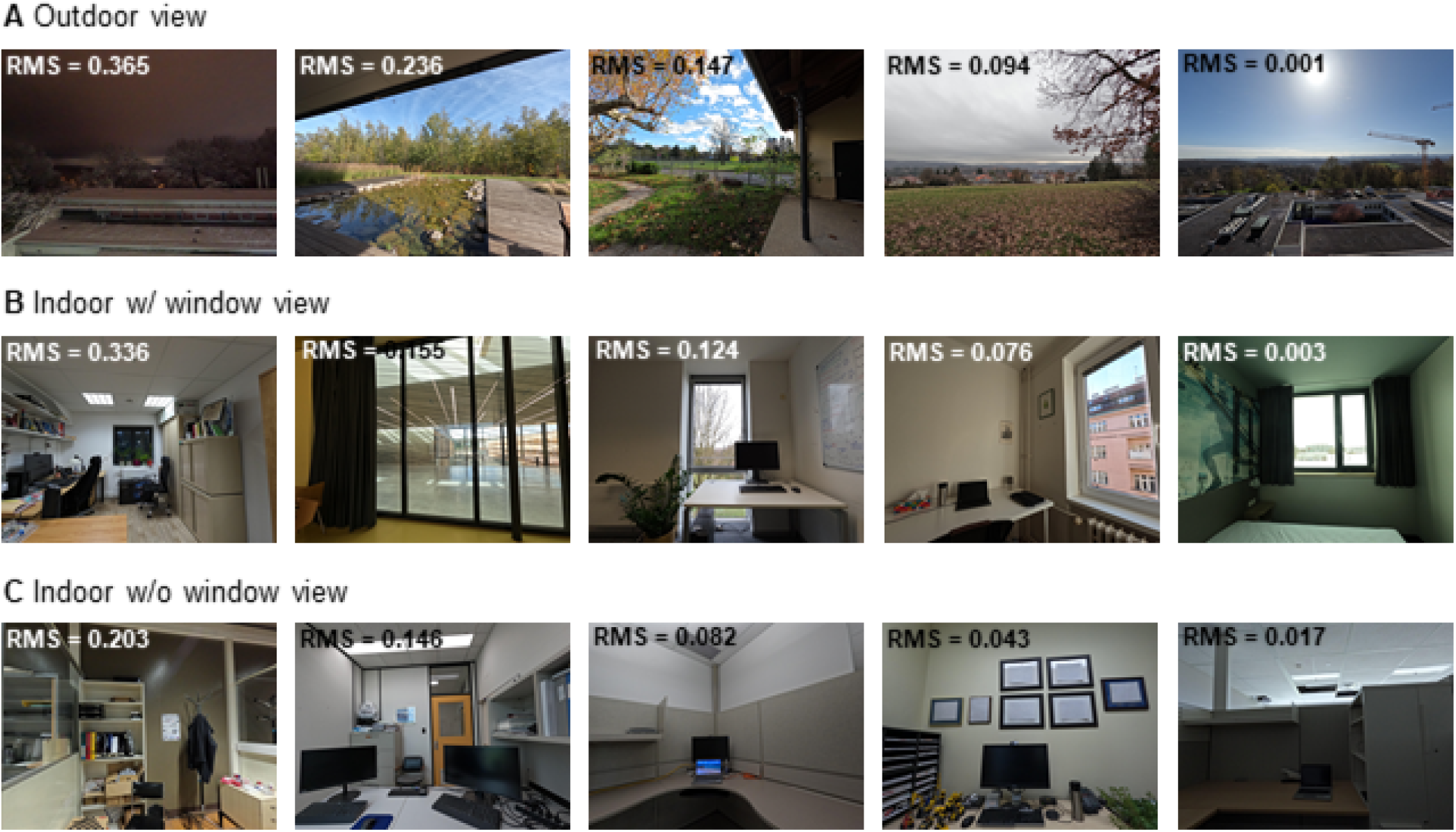
Example RGB images illustrate the range of melanopic RMS contrast observed across view categories. A.) Outdoor-view images ordered from highest to lowest melanopic RMS contrast, ranging from 0.365 to 0.001. B.) Indoor-with-window-view images ordered from highest to lowest melanopic RMS contrast, ranging from 0.336 to 0.003. C.) Indoor-without-window-view images ordered from highest to lowest melanopic RMS contrast, ranging from 0.203 to 0.017. RMS contrast values are image-level values calculated from the corresponding *α*-opic radiance maps.

### Spatial statistics of natural scenes

Natural scenes show consistent spatial structure, often following a power-law distribution in their amplitude spectra^20,42,44^. This means that amplitude decreases systematically with increasing spatial frequency. Such patterns have been well documented in luminance and cone-based images and are thought to reflect visual system adaptation. Most previous studies focused on cone responses under daylight conditions. In contrast, the spatial statistics of rod and melanopsin channels have received little attention. Here, we analyse all five *α*-opic channels and composite chromatic channels (L+M+S, L-M, S-(L+M)) using direct multispectral image acquisition without relying on spectral modelling. This allows a more complete and accurate view of visual environments as the human retina encodes them.

Figure 6 presents two example scenes illustrating the range of spatial frequency slopes observed in the melanopic channel. Scene A shows the flattest slope (*−*0.20), indicating weak spatial modulation and relatively uniform melanopic content in all directions. Scene B has the steepest slope (*−*1.74), reflecting a dominance of low-frequency structure and strong spatial variation. In both cases, the melanopic amplitude spectra are highly correlated with spatial frequency (*r* = *−*0.828 and *r* = *−*0.993, *p <* 10*^−^*^5^), confirming that a consistent power-law relationship is preserved despite variation in slope. Notably, both the flattest and steepest slopes originate from the outdoor view category. This indicates that outdoor scenes show greater variability in spatial frequency and, consequently, in their spatial structure compared with the more homogeneous indoor categories.

**Figure 6:**
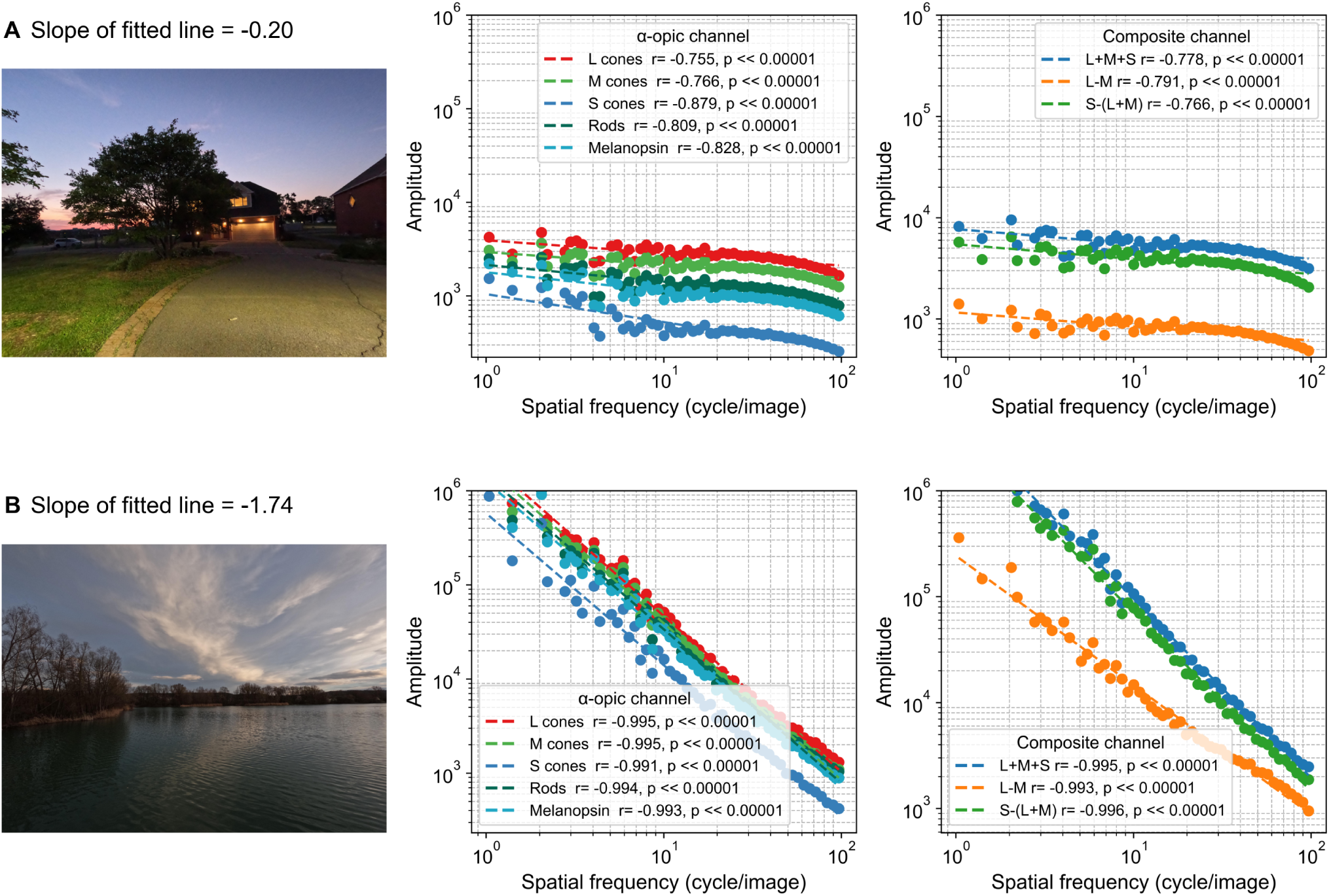
Example scenes illustrate the range of spatial-frequency slopes in melanopic amplitude spectra. A.) Scene with the least steep decline in the melanopic amplitude spectrum. The left panel shows the corresponding wide-angle RGB image, the middle panel shows amplitude as a function of spatial frequency for the five *α*-opic channels, and the right panel shows amplitude as a function of spatial frequency for composite channels derived from the *α*-opic channels. Lines represent linear fits to log-transformed amplitude spectra, and the reported slope describes the decline in amplitude with increasing spatial frequency. B.) Scene with the steepest negative melanopic amplitude-spectrum slope, shown using the same layout and measures as in panel A.

To examine how spatial frequency content varies across environments, we plotted the probability density distribution of the slopes fitted to the log-log amplitude spectra of the melanopic channel for each view category, as shown in Figure 7. The slope values range from approximately *−*1.8 to *−*0.2, consistent with the power-law distribution observed in natural scenes in previous research. However, the slope distributions differ clearly between indoor and outdoor conditions.

**Figure 7:**
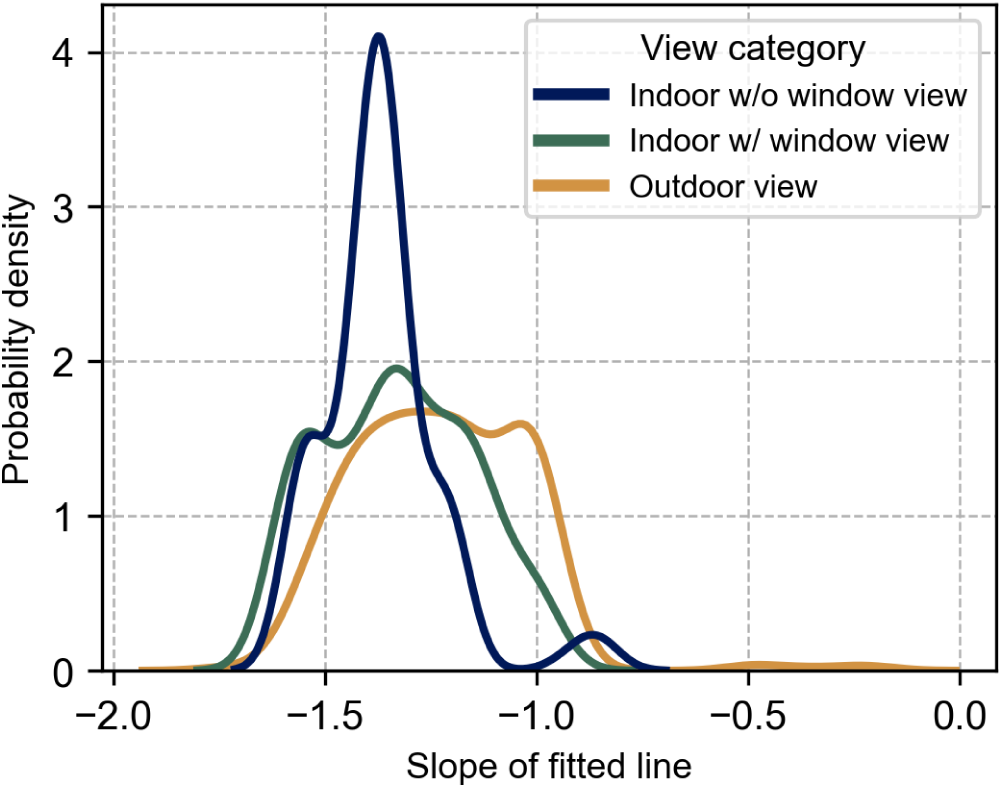
Melanopic amplitude-spectrum slopes differ across indoor and outdoor view categories. Probability density plot showing the distribution of slopes from linear fits to log-transformed melanopic amplitude spectra, grouped by view category. Density curves represent smoothed probability densities of image-level slope values, not means or error bars.

Indoor scenes without a window view exhibit the steepest slopes, clustering around *−*1.4 to *−*1.3, indicating a dominance of low spatial frequencies and relatively uniform spatial structure in melanopic contrast. Indoor scenes with window views show a slightly flatter distribution, with slopes more broadly distributed and centred around *−*1.3 to *−*1.2. Outdoor scenes have the flattest slopes, generally between *−*1.2 and *−*1.0, suggesting a higher contribution of fine spatial details and a broader range of spatial frequencies in these environments.

These results align with previous studies on the spatial statistics of natural scenes^1,15,19,42,45–47^, supporting the notion that 1/f-like spectral characteristics are preserved even in photoreceptor-specific representations such as the melanopic channel.

## Discussion

This study provides a comprehensive characterisation of the spatial and spectral statistics of natural scenes across a diverse dataset sampled from indoor and outdoor environments. Using image- and spectral-based analyses, we quantified contrast, amplitude spectra, and cross-photoreceptor correlations for all five human photoreceptors (L, M, S cones, rods, and melanopsin), thereby describing the structure of environmental light in terms of *α*-opic signals. While previous research has extensively examined natural image statistics in relation to cone-mediated vision^1,19,46,48^, little attention has been given to how these spectral and spatial properties extend to melanopic channels^40,41,49,50^. By characterising these distributions across ecological contexts, our results provide critical data for future models linking environmental image statistics to photoreceptor-specific adaptation and visual, non-visual processing.

Radiance and irradiance levels vary systematically across view types. Indoor environments without windows show consistently low values across all channels, reflecting uniform electric lighting. Indoor scenes with window views exhibit intermediate values, influenced by varying daylight entry. Outdoor scenes show the highest radiance and irradiance, particularly for M cones, which aligns with the spectral profile of daylight. These findings are consistent with previous daylight studies but extend them by including all five *α*-opic channels.

Photopic and melanopic illuminance values are strongly correlated across all scenes (*r ≈* 0.9*, p <* 10*^−^*^5^), regardless of view type or time of day. A moderate-to-strong correlation (*r ≈* 0.7*, p <* 10*^−^*^5^) is also observed between spectral melanopic EDI and image-based melanopic luminance, with outdoor scenes showing the strongest alignment. This supports the reliability of image-based metrics for estimating photoreceptor-relevant light exposure, especially in natural settings. It is important to note that, given the strong correlations between melanopic and cone-mediated radiance across environments, the spatial statistics reported here are best interpreted as reflecting the structure of the environmental light field rather than evidence for melanopsin-specific spatial encoding.

Outdoor scenes exhibit greater variability in RMS contrast compared to indoor environments. Higher melanopic contrast is associated with increased melanopic luminance in outdoor and windowed indoor scenes. No such association is found in windowless rooms, likely due to uniform lighting. These patterns agree with prior reports of contrast structure in natural images^16^, and extend them to melanopic and rod-based contrast.

The amplitude spectra of melanopic images follow a power-law distribution with slopes between *−*0.2 and *−*1.8, consistent with 1/f statistics observed in cone-based studies^45^. Steeper slopes in indoor scenes suggest reduced high-frequency structure, while flatter slopes in outdoor views reflect richer spatial detail. This confirms that scale-invariant (1/f-like) properties of natural scenes extend to all photoreceptor channels, including melanopsin and rods, which have been underexplored in previous work; given the strong inter-channel correlations and the fact that edges and luminance transitions occur across multiple spatial scales in natural environments, these melanopic 1/f statistics are most parsimoniously explained by shared environmental structure rather than a melanopsin-specific spatial processing mechanism.

This work provides both a new photoreceptor-resolved natural-scene dataset and a systematic characterisation of its spectral and spatial statistics. Across scene categories, we identify several robust descriptive patterns, including higher *α*-opic exposure outdoors than indoors, strong coupling between melanopic and photopic measures, greater contrast variability in outdoor scenes, and 1/f-like amplitude spectra with scene-dependent slopes. We anticipate that this resource will support future experimental, computational, and ecological studies of visual and non-visual photoreception.

## Limitations of the Study

The technical limitations of this study include a small parallax due to the measurement devices not being co-planar, which is inevitable without a conjoint radiance/irradiance measurement setup. However, we believe these minor differences are not consequential for the main conclusions of the manuscript. Temporal uncertainty arises from different integration times, which are driven by practical demands due to the varying range of possible light levels. The field of view of the circadiometer camera is limited and does not exactly match the cosine-corrected field of view of our spectral irradiance measurements.

Regarding variability in natural scenes, our sampling was structured around predefined scene categories and diverse accessible locations, but it was not intended to constitute an exhaustive or population-representative sample of all possible natural scenes. While the dataset spans multiple indoor and outdoor subcategories across five geographic locations, it still represents a subset of possible real-world environments, and broader future sampling would further improve coverage and comprehensiveness.

Although some measurements were obtained from the same scene geometries, the substantial variation in illumination conditions and corresponding *α*-opic radiance distributions resulted in distinct spatial contrast and spectral characteristics; however, a limited degree of structural dependence due to shared geometry cannot be entirely excluded. As such, the dataset represents an opportunity for analyses of illumination changes while scene geometry is kept constant.

An additional limitation is that our analyses characterise the environmental light field as captured by the measurement devices, rather than the retinal image after transformation by the optics of the eye. In the present work, we did not model optical factors such as ocular blur, aberrations, scatter, or other spatial distortions, nor did we account for retinal sampling by the photoreceptor mosaic. Accordingly, the spatial statistics reported here should be interpreted as properties of the scene-side signal available to the visual system, not as a full account of how that signal is represented at the retinal level. Incorporating ocular optics and photoreceptor sampling will be an important direction for future work, as these factors will introduce additional degradations and filtering before the signal reaches the retina^41,51^.

Future studies could explore more detailed categories beyond different views to better understand natural scenes. Comprehensive metadata makes it possible to examine subcategories within scenes with specific spatial or lighting characteristics.

## Resource availability

### Lead contact

Requests for further information and resources should be directed to and will be fulfilled by the lead contact, Manuel Spitschan.

### Materials availability

This study did not generate new materials.

### Data and code availability

- The SCENES raw dataset is deposited in the EDMOND general data repository under the name *SCENES* (https://doi.org/10.17617/3.PYHUO5). The derived NumPy arrays generated from the original radiance maps, along with tabular aggregated spatial metrics, are stored in a second repository named *SCENES derivatives* (https://doi.org/10.17617/3.NX2H2U).
- All code has been deposited as a GitHub repository (https://github.com/tscnlab/Tabande iScience_2026) archived on Zenodo (https://doi.org/10.5281/zenodo.19434780). We used the *pearsonr* function from the *SciPy* Python library (version 1.16.1)^52^ to calculate the Pearson correlation coefficient and the associated p-value for testing the null hypothesis of no correlation between two metrics. The correlation coefficients were calculated using the *corr* function from the Pandas Python library^53,54^, which computes the pairwise Pearson correlation coefficient. This function generates correlation matrices for the five *α*-opic metrics. We used the *fft*, *fftshift*, and *linregress* functions from the *SciPy* library (version 1.16.1) in Python to compute amplitude spectra.
- Any additional information required to reanalyse the data reported in this paper is available from the lead contact upon request.

## Acknowledgments

We sincerely thank all external collaborators and members of the Translational Sensory & Circadian Neuroscience Unit (MPS/TUM/TUMCREATE) for their support, as well as all contributors to the dataset.

## Author contributions

Conceptualisation: N.T., M.S.

Data curation: N.T.

Formal analysis: N.T.

Funding acquisition: M.S.

Methodology: N.T., M.S.

Resources: M.S.

Supervision: M.S.

Visualisation: N.T.

Writing – original draft: N.T.

Writing – review & editing: N.T., M.S.

## Declaration of interests

N.T. declares no conflicts of interest.

M.S. declares the following potential conflicts of interest in the past five years (2021–2025). *Academic roles*: Member of the Board of Directors, Society of Light, Rhythms, and Circadian Health (SLRCH); Chair of Joint Technical Committee 20 (JTC20) of the International Commission on Illumination (CIE); Member of the Daylight Academy; Chair of Research Data Alliance Working Group Optical Radiation and Visual Experience Data. *Remunerated roles*: Speaker of the Steering Committee of the Daylight Academy; Ad-hoc reviewer for the Health and Digital Executive Agency of the European Commission; Ad-hoc reviewer for the Swedish Research Council; Associate Editor for LEUKOS, journal of the Illuminating Engineering Society; Examiner, University of Manchester; Examiner, Flinders University; Examiner, University of Southern Norway. *Funding*: Received research funding and support from the Max Planck Society, Max Planck Foundation, Max Planck Innovation, Technical University of Munich, Wellcome Trust, National Research Foundation Singapore, European Partnership on Metrology, VELUX Foundation, Bayerisch-Tschechische Hochschulagentur (BTHA), BayFrance (Bayerisch-Franzö sisches Hochschulzentrum), BayFOR (Bayerische Forschungsallianz), and Reality Labs Research. Honoraria for talks: Received honoraria from the ISGlobal, Research Foundation of the City University of New York and the Stadt Ebersberg, Museum Wald und Umwelt. *Travel reimbursements*: Daimler und Benz Stiftung. *Patents*: Named on European Patent Application EP23159999.4A (“System and method for corneal-plane physiologically-relevant light logging with an application to personalised light interventions related to health and well-being”). With the exception of the funding source supporting this work, M.S. declares no influence of the disclosed roles or relationships on the work presented herein. The funders had no role in study design, data collection and analysis, decision to publish or preparation of the manuscript.

## Supplemental information index

Figures S1 related to ”A comprehensive *α*-opic dataset of natural scenes” subsection in a PDF

## STAR methods

### Method details

#### Dataset and data collection

We conducted a comprehensive data collection campaign to characterise the spectral, spatial, and temporal properties of natural scenes. The multimodal measurement setup comprises an *α*-opic imaging circadiometer, a high-resolution spectroradiometer, an uncalibrated wide-field RGB camera, and a temperature and humidity logger. All instruments were integrated into a portable box (42 *×* 32 *×* 27 cm). This custom measurement box was mounted on a tripod. The equipment was powered by external batteries or a direct power source. This setup was suitable for indoor and outdoor measurements throughout the day. Data were collected across various times of day, seasons, and different geographical locations. In each measurement bin, we collected data from all measurement devices and described the scenes using a newly developed metadata descriptor (n=43 items), encompassing detailed information including geographical information, weather conditions, and scene categories. Figure 1A shows the setup and measurement procedure diagram in each measurement bin. Repeated measurements were prioritised for scenes that were accessible across multiple times of day and expected to show meaningful daylight-driven variation.

For the spatial analysis, we used spectral data from the spectroradiometer and *α*-opic radiance maps from the circadiometer.

- *Imaging circadiometer* : The Imaging Circadiometer (WP690E-Circ, Westboro Photonics, Ottawa, Canada; 14 mm, 40*^◦^×* 48*^◦^*, 9.1 megapixels, 4 *×* 10*^−^*^6^ to 60 W*/*m^2^*/*sr) is equipped with five in-built filters, corresponding to L-, M-, S-, rod-, and melanopic human spectral sensitivities. During each capture, the camera’s motorised system rotates these filters sequentially, producing five distinct *α*-opic images from the scene. The camera is equipped with automated electronic lens control and captures high dynamic range (HDR) images for each *α*-opic channel. Each radiance map has dimensions of 2712 *×* 3388 pixels.
- *Spectroradiometer* : We utilised the high-resolution spectroradiometer (Spectraval 1511 HiRes, Jena Technische Instrumente GmbH, Jena, Germany; 380-780 nm, 1 nm resolution, and spectral luminance ranging from 0.02 to 18*×* 10^4^ cd*/*m^2^). We used a cosine diffuser to measure the spectral illuminance of the scenes. The spectroradiometer derives numerous metrics from spectral illuminance. This analysis focused on melanopic equivalent daylight illuminance (melanopic EDI), photopic illuminance, and the five *α*-opic irradiances.

#### Image analysis

The circadiometer camera outputs five *α*-opic radiance maps. We used the device’s proprietary software (Photometrica) https://wphotonics.com/software/ and stored them in .pmm format. Using the same software for each measurement, these five channels were exported along with six derived channels (luminance, trix, triz, stdred, stdgreen, stdblue) as an 11-channel dataset. This dataset was then saved as an ND NumPy array within a .npz format for further analysis. In the quality control process, we encountered two types of invalid pixels, represented as NaN values and infinite (Inf) values in ND NumPy arrays. Missing pixels (NaN values) occur due to misalignment between filters and the lens. The number of missing pixels in each channel was fixed and did not change across different measurements.

Inf values in ND NumPy arrays correspond to overexposed pixels. Overexposed pixels arise due to the technical limitations of the camera’s CCD sensor. The sensor has a maximum radiance threshold. When the light intensity surpasses this limit, the sensor cannot accurately record the values, resulting in the pixel values being represented as infinite (Inf) in the ND NumPy arrays. In the next step, we calculated the average radiance in each channel. This parameter is the mean value of all valid pixels in each radiance map. We also computed the mean luminance for each *α*-opic channel using the conversion factor for each human photoreceptor^55^.

Subsequent spatial analyses were performed using the five *α*-opic NumPy arrays. The first analysis involved calculating the root mean square (RMS) contrast, a metric that quantifies the global variation in pixel intensity across an image. The second analysis computed the amplitude spectra to assess the spatial frequency characteristics of the images.

#### Root mean square (RMS) contrast

While contrast can be quantified using several metrics^56,57^ (e.g., Michelson and Weber contrast), Bex and Makous^42^ demonstrated that RMS contrast is a useful contrast indicator for natural scenes, reflecting overall luminance variability. The RMS contrast quantifies the local variation in pixel intensity across an image^58^.

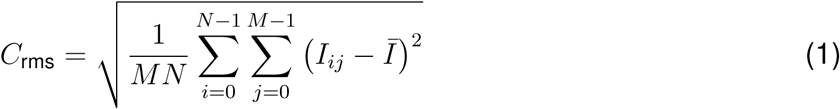

In Equation (1), M and N are the dimensions of the image, *I_ij_*represents the intensity of the (i,j) pixel, and *I̅* is the mean intensity of the image. In the RMS contrast calculation, we normalised the pixel values to the range [0, 1] by dividing by the maximum finite value in the image^42^. Non-finite pixel values (e.g., NaN or Inf) are excluded from the calculation. That is, the RMS contrast is computed as the square root of the average squared deviation of the normalised pixel intensities from their mean.

We calculated the global RMS contrast of five *α*-opic channels for each image in the dataset.

#### Amplitude spectra

The amplitude spectrum analysis quantifies the spatial frequency content of an image^20,59^ by following these steps:

- **Computing the amplitude spectrum:** First, we cropped the input image to a centred square patch, using the smaller dimension (height or width) to keep it uniform. To fix irregularities, we replaced NaN values with the median of valid pixels and substituted Inf values with the maximum valid pixel. Next, we applied a 2D Fast Fourier Transform (FFT) to convert the image into the frequency domain, shifting the zero-frequency component to the centre. Finally, we extracted the amplitude spectrum by computing the magnitude of the FFT, which captures the intensity of different spatial frequencies in the image.
- **Radial frequency distribution:** We analysed the amplitude spectrum by dividing it into radial frequency bins, which are logarithmically spaced to cover a broad range of spatial frequencies. We then calculated the radial distances from the centre of the spectrum and aggregated the amplitude values within each bin. We normalised the radial amplitude by the number of contributing pixels per bin to obtain an average amplitude for each frequency. Frequencies with near-zero amplitudes and those exceeding 10^2^ are excluded to focus on the meaningful spatial frequency content^20^. This process captures the spatial frequency distribution and radial variation of each image.
- **Least-squares linear regression for amplitude spectra:** For each measurement, we calculated the amplitude spectra for the five *α*-opic channels (original image channels) as well as their opponent colour channels (*L* + *M* + *S*, *L − M*, *S −* (*L* + *M*)). We then transformed the radial frequencies and amplitudes from the amplitude spectrum analysis to a logarithmic scale to linearise the relationship between these variables. Finally, we performed linear regression on the log-transformed data, which yields a slope and offset that describe the spectral characteristics of each image.

The fitted line is computed from the regression equation:

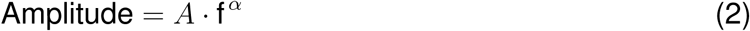

where *A* represents the offset, a constant that scales the amplitude, and *α* is the slope, determining how the amplitude changes with frequency.

### Quantification and statistical analysis

We calculated descriptive statistics, including the mean, standard deviation, median, minimum, maximum, and interquartile range (IQR), for the following variables: melanopic equivalent daylight illuminance (melanopic EDI), photopic illuminance, the five *α*-opic irradiances from the spectroradiometer, and the mean *α*-opic radiances from the circadiometer images.

We used the *pearsonr* function from the *SciPy* Python library (version 1.16.1)^52^ to calculate the Pearson correlation coefficient and the associated p-value for testing the null hypothesis of no correlation between two metrics. The correlation coefficients were calculated using the *corr* function from the Pandas Python library^53,54^, which computes the pairwise Pearson correlation coefficient. This function generates correlation matrices for the five *α*-opic metrics. We used the *fft*, *fftshift*, and *linregress* functions from the *SciPy* library (version 1.16.1) in Python to compute amplitude spectra.

## Notes

https://doi.org/10.17617/3.PYHUO5

https://doi.org/10.17617/3.NX2H2U

https://github.com/tscnlab/TabandehEtAl_iScience_2026

https://doi.org/10.5281/zenodo.19434780

### Summary of Updates

Summary and section headings updated for journal style; dataset count corrected from 675 to 671 measurements; Discussion and limitations clarified; Resource availability and STAR methods added; figure legends revised; supplemental timelapse figure and supplemental information index added.

## References

1. Geisler, W.S. (2008). Visual perception and the statistical properties of natural scenes. Annual Review of Psychology 59, 167–192. doi: 10.1146/annurev.psych.58.110405.085632.

2. Tkačik, G., Garrigan, P., Ratliff, C., Milčinski, G., Klein, J.M., Seyfarth, L.H., Sterling, P., Brainard, D.H., and Balasubramanian, V. (2011). Natural images from the birthplace of the human eye. PLOS ONE 6, e20409. doi: 10.1371/journal.pone.0020409.

3. Caves, E.M., Cheney, K.L., Dacke, M., Dixit, T., Fialko, K., Franklin, A.M., Jessop, A.L., Hart, N.S., Hempel de Ibarra, N., Morehouse, N.I., Morgan, R., Murugavel, B., Oakley, T.H., Speiser, D.I., Stoddard, M.C., Warrant, E.J., Johnsen, S., and Schweikert, L.E. (2025). Emerging frontiers in visual ecology. Journal of Experimental Biology 228, jeb250537. doi: 10.1242/jeb.250537.

4. Simoncelli, E.P., and Olshausen, B.A. (2001). Natural image statistics and neural representation. Annual Review of Neuroscience 24, 1193–1216. doi: 10.1146/annurev.neuro.24.1.1193.

5. Marcos, S. (2025). Optical and visual diet in myopia. Investigative Ophthalmology and Visual Science 66, 3. doi: 10.1167/iovs.66.7.3.

6. Webster, M.A., MacLin, O.H., Rees, A.L., and Raker, V.E. (1996). Contrast adaptation and the spatial structure of natural images. Perception 25, 174–174. doi: 10.1068/v96l1011.

7. Tolhurst, D.J., and Tadmor, Y. (1997). Discrimination of changes in the slopes of the amplitude spectra of natural images: Band-limited contrast and psychometric functions. Perception 26, 1011–1025. doi: 10.1068/p261011.

8. Párraga, C.A., and Tolhurst, D.J. (2000). The effect of contrast randomisation on the discrimination of changes in the slopes of the amplitude spectra of natural scenes. Perception 29, 1101–1116. doi: 10.1068/p2904.

9. Hansen, T., and Gegenfurtner, K.R. (2009). Independence of color and luminance edges in natural scenes. Visual Neuroscience 26, 35–49. doi: 10.1017/S0952523808080796.

10. Bex, P. (2007). Contrast gain control in natural scenes. Journal of Vision. URL: https://www.academia.edu/109210213/Contrast_gain_control_in_natural_scenes.

11. Foster, D.H., Amano, K., Nascimento, S.M.C., and Foster, M.J. (2006). Frequency of metamerism in natural scenes. JOSA A 23, 2359–2372. doi: 10.1364/JOSAA.23.002359.

12. Bex, P.J., Solomon, S.G., and Dakin, S.C. (2009). Contrast sensitivity in natural scenes depends on edge as well as spatial frequency structure. Journal of Vision 9, 1. doi: 10.1167/9.10.1.

13. Jarvis, J., Triantaphillidou, S., and Gupta, G. (2022). Contrast discrimination in images of natural scenes. JOSA A 39, B50–B64. doi: 10.1364/JOSAA.447390.

14. Triantaphillidou, S., Jarvis, J., Psarrou, A., and Gupta, G. (2019). Contrast sensitivity in images of natural scenes. Signal Processing: Image Communication 75, 64–75. doi: 10.1016/j.image.2019.03.002.

15. Rideaux, R., West, R.K., Wallis, T.S.A., Bex, P.J., Mattingley, J.B., and Harrison, W.J. (2022). Spatial structure, phase, and the contrast of natural images. Journal of Vision 22, 4. doi: 10.1167/jov.22.1.4.

16. Frazor, R.A., and Geisler, W.S. (2006). Local luminance and contrast in natural images. Vision Research 46, 1585–1598. doi: 10.1016/j.visres.2005.06.038.

17. Tolhurst, D.J., Tadmor, Y., and Arthurs, G. (1996). Detection of changes in the amplitude spectra of natural images is explained by a band-limited local-contrast model. In Human Vision and Electronic Imaging vol. 2657. SPIE pp. 154–165. doi: 10.1117/12.238712.

18. Ruderman, D.L., and Bialek, W. (1994). Statistics of natural images: Scaling in the woods. Physical Review Letters 73, 814–817. doi: 10.1103/PhysRevLett.73.814.

19. Burton, G.J., and Moorhead, I.R. (1987). Color and spatial structure in natural scenes. Applied Optics 26, 157–170. doi: 10.1364/AO.26.000157.

20. Tolhurst, D.J., Tadmor, Y., and Chao, T. (1992). Amplitude spectra of natural images. Ophthalmic and Physiological Optics 12, 229–232. doi: 10.1111/j.1475-1313.1992.tb00296.x.

21. Brady, N., and Field, D.J. (2000). Local contrast in natural images: Normalisation and coding efficiency. Perception 29, 1041–1055. doi: 10.1068/p2996.

22. Webster, M.A., and Mollon, J.D. (1997). Adaptation and the color statistics of natural images. Vision Research 37, 3283–3298. doi: 10.1016/s0042-6989(97)00125-9.

23. Mure, L.S., Vinberg, F., Hanneken, A., and Panda, S. (2019). Functional diversity of human intrinsically photosensitive retinal ganglion cells. Science 366, 1251–1255. doi: 10.1126/science.aaz0898.

24. Schlangen, L.J.M., and Price, L.L.A. (2021). The lighting environment, its metrology, and non-visual responses. Frontiers in Neurology 12, 624861. doi: 10.3389/fneur.2021.624861.

25. Blume, C., Garbazza, C., and Spitschan, M. (2019). Effects of light on human circadian rhythms, sleep and mood. Somnologie 23, 147–156. doi: 10.1007/s11818-019-00215-x.

26. Nowozin, C., Wahnschaffe, A., Rodenbeck, A., de Zeeuw, J., Hä del, S., Kozakov, R., Schö pp, H., Mü nch, M., and Kunz, D. (2017). Applying melanopic lux to measure biolog-ical light effects on melatonin suppression and subjective sleepiness. Current Alzheimer Research 14, 1042–1052. doi: 10.2174/1567205014666170523094526.

27. Prayag, A.S., Najjar, R.P., St Hilaire, M.A., Rahman, S.A., Rajaratnam, S.M.W., Rü ger, M., Brainard, G.C., Czeisler, C.A., Andersen, M., Gooley, J.J., and Lockley, S.W. (2019). Melatonin suppression is exquisitely sensitive to light and primarily driven by melanopsin in humans. Journal of Pineal Research 66, e12562. doi: 10.1111/jpi.12562.

28. Didikoglu, A., Mohammadian, N., Johnson, S., van Tongeren, M., Wright, P., Casson, A.J., Brown, T.M., and Lucas, R.J. (2023). Associations between light exposure and sleep timing and sleepiness while awake in a sample of uk adults in everyday life. Proceedings of the National Academy of Sciences 120, e2301608120. doi: 10.1073/pnas.2301608120.

29. Lucas, R.J., Peirson, S., Berson, D.M., Brown, T.M., Cooper, H.M., Czeisler, C.A., Figueiro, M.G., Gamlin, P.D., Lockley, S.W., O’Hagan, J.B., Price, L.L.A., Provencio, I., Skene, D.J., and Brainard, G.C. (2014). Measuring and using light in the melanopsin age. Trends in Neurosciences 37, 1–9. doi: 10.1016/j.tins.2013.10.004.

30. Allen, A.E., Martial, F.P., and Lucas, R.J. (2017). Melanopsin contributions to the representation of images in the early visual system. Current Biology 27, 1623–1632.e4. doi: 10.1016/j.cub.2017.04.046.

31. Mure, L.S. (2021). Intrinsically photosensitive retinal ganglion cells of the human retina. Frontiers in Neurology 12, 636330. doi: 10.3389/fneur.2021.636330.

32. Spitschan, M. (2019). Melanopsin contributions to non-visual and visual function. Current Opinion in Behavioral Sciences 30, 67–72. doi: 10.1016/j.cobeha.2019.06.004.

33. Do, M.T., Kang, S.H., Xue, T., Zhong, H., Liao, H.W., Bergles, D.E., and Yau, K.F. (2009). Photon capture and signalling by melanopsin retinal ganglion cells. Nature 457, 281–287. doi: 10.1038/nature07682.

34. Berson, D.M., Dunn, F.A., and Takao, Y. (2002). Intrinsically photosensitive retinal ganglion cells and the pupillary light reflex. Science 295, 1071–1073. doi: 10.1126/science.1067262.

35. Price, L.L.A., and Blattner, P. (2022). Circadian and visual photometry. Progress in Brain Research 273, 1–11. doi: 10.1016/bs.pbr.2022.02.014.

36. Nugent, T.W., and Zele, A.J. (2025). What can the eye see with melanopsin? Proceedings of the National Academy of Sciences. doi: 10.1073/pnas.2411151121.

37. Brown, T.M., Brainard, G.C., Cajochen, C., Czeisler, C.A., Hanifin, J.P., Lockley, S.W., Lucas, R.J., Mü nch, M., O’Hagan, J.B., Peirson, S.N., Price, L.L.A., Roenneberg, T., Schlangen, L.J.M., Skene, D.J., Spitschan, M., Vetter, C., Zee, P.C., and Wright, K.P. (2022). Recommendations for daytime, evening, and nighttime indoor light exposure to best support physiology, sleep, and wakefulness in healthy adults. PLOS Biology 20. doi: 10.1371/journal.pbio.3001571.

38. Webler, F.S., Spitschan, M., Foster, R.G., Andersen, M., and Peirson, S.N. (2019). What is the ‘spectral diet’ of humans? Current Opinion in Behavioral Sciences 30, 80–86. doi: 10.1016/j.cobeha.2019.06.006.

39. Brown, T.M. (2020). Melanopic illuminance defines the magnitude of human circadian light responses under a wide range of conditions. Journal of Pineal Research 69, e12655. doi: 10.1111/jpi.12655.

40. Spitschan, M., and Aguirre, G.K. (2017). Vision: Melanopsin as a raumgeber. Current Biology 27, R644–R646. doi: 10.1016/j.cub.2017.05.052.

41. Barrionuevo, P.A., and Diaz-Barrancas, F. (2025). Melanopsin-mediated image statistics from natural and human-made environments. Scientific Reports 15, 29965. doi: 10.1038/s41598-025-15981-y.

42. Bex, P.J., and Makous, W. (2002). Spatial frequency, phase, and the contrast of natural images. JOSA A 19, 1096–1106. doi: 10.1364/JOSAA.19.001096.

43. Jakhetiya, V., Lin, W., Jaiswal, S., Gu, K., and Guntuku, S.C. (2018). Just noticeable difference for natural images using rms contrast and feedback mechanism. Neurocomputing 275, 366–376. doi: 10.1016/j.neucom.2017.08.031.

44. Field, D.J. (1994). What is the goal of sensory coding? Neural Computation 6, 559–601. doi: 10.1162/neco.1994.6.4.559.

45. Ruderman, D.L. (1994). The statistics of natural images. Network: Computation in Neural Systems 5, 517. doi: 10.1088/0954-898X/5/4/006.

46. Ruderman, D.L., Cronin, T.W., and Chiao, C.C. (1998). Statistics of cone responses to natural images: implications for visual coding. JOSA A 15, 2036–2045. doi: 10.1364/JOSAA.15.002036.

47. Dyakova, O., Lee, Y.J., Longden, K.D., Kiselev, V.G., and Nordströ m, K. (2015). A higher order visual neuron tuned to the spatial amplitude spectra of natural scenes. Nature Communications 6, 8522. doi: 10.1038/ncomms9522.

48. Shevell, S.K., and Kingdom, F.A.A. (2008). Color in complex scenes. Annual Review of Psychology 59, 143–166. doi: 10.1146/annurev.psych.59.103006.093619.

49. Allen, A.E., Martial, F.P., and Lucas, R.J. (2019). Form vision from melanopsin in humans. Nature Communications 10, 2274. doi: 10.1038/s41467-019-10113-3.

50. Ponting, S., Waskett, R.K., Spitschan, M., and Smithson, H.E. (2025). Quantifying cie alpha-opic signals in the indoor built environment. J. Opt. Soc. Am. A 42, B379–B390. doi: 10.1364/JOSAA.545151.

51. Wandell, B.A., Brainard, D.H., and Cottaris, N.P. (2022). Visual encoding: Principles and software. Progress in Brain Research 273, 199–229. doi: 10.1016/bs.pbr.2022.04.006.

52. Virtanen, P., Gommers, R., Oliphant, T.E., Haberland, M., Reddy, T., Cournapeau, D. et al. (2020). Scipy 1.0: Fundamental algorithms for scientific computing in python. Nature Methods 17, 261–272. doi: 10.1038/s41592-019-0686-2.

53. McKinney, W. (2010). Data structures for statistical computing in python. In van der S. Walt, and J. Millman, eds. Proceedings of the 9th Python in Science Conference. pp. 56–61. doi: 10.25080/Majora-92bf1922-00a.

54. The pandas development team (2020). pandas-dev/pandas: Pandas. Zenodo. URL: https://doi.org/10.5281/zenodo.3509134. doi: 10.5281/zenodo.3509134 version latest.

55. Spitschan, M., Mead, J., Roos, C., Lowis, C., Griffiths, B., Mucur, P., Herf, M., Nam, S., and Veitch, J.A. (2022). luox: validated reference open-access and open-source web platform for calculating and sharing physiologically relevant quantities for light and lighting. Wellcome Open Research 6, 69. doi: 10.12688/wellcomeopenres.16595.3.

56. Kukkonen, H., Rovamo, J., Tiippana, K., and Nä sä nen, R. (1993). Michelson contrast, rms contrast and energy of various spatial stimuli at threshold. Vision Research 33, 1431–1436. doi: 10.1016/0042-6989(93)90049-3.

57. Pelli, D.G., and Bex, P. (2013). Measuring contrast sensitivity. Vision research 90, 10–14. doi: 10.1016/j.visres.2013.04.015.

58. Peli, E. (1990). Contrast in complex images. JOSA A 7, 2032–2040. doi: 10.1364/JOSAA.7.002032.

59. Hansen, B., and Essock, E. (2005). Influence of scale and orientation on the visual perception of natural scenes. Visual Cognition 12, 1199–1234. doi: 10.1080/13506280444000715.

