## Supplementary Materials for "Photoreceptor-specific scene statistics reveal melanopic structure in natural environments"

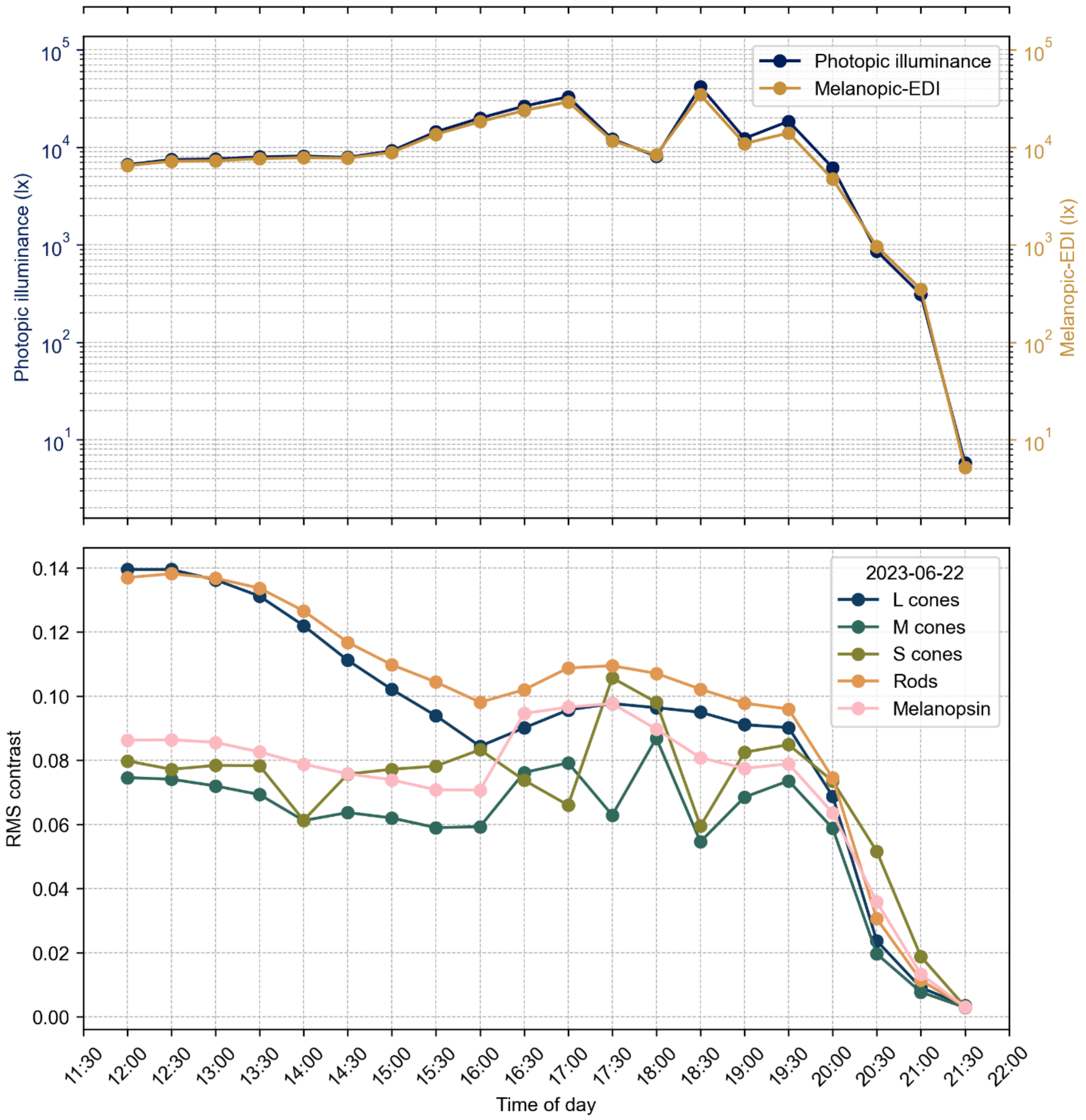


**Figure S1. Example timelapse of a natural scene illustrating temporal changes in α-opic metrics and spatial contrast**

Spatiotemporal variation of a sample timelapse on 2023-06-22: example scene images across the day (top), corresponding photopic illuminance and melanopic EDI (log scale, middle), and photoreceptor-specific RMS contrast (bottom), showing dynamic changes in repeated measurements across a day.
